# D-LMBmapX: Generalised Deep Learning Pipeline for 5D Whole-brain Circuitry Profiling

**DOI:** 10.1101/2025.02.25.639766

**Authors:** Zhongyu Li, Peiqi Li, Hangzhen Pan, Annabel M.J. Adams, Zuqi Huang, Yaoyu Su, He Liu, Jinjiang Li, Baoming Zhong, Yuqin Su, Liam Bray, Robyn N. Sturgess, Chen Li, Jihua Zhu, Dingwen Zhang, Shaoting Zhang, Jing Ren

**Affiliations:** Neurobiology Division, MRC Laboratory of Molecular Biology, Francis Crick Avenue, Cambridge, CB2 0QH, United Kingdom; School of Computer Science, Shanghai Jiao Tong University, Shanghai, China; School of Software Engineering, Xi’an Jiaotong University, Xi’an, China; Hefei Comprehensive National Science Center, Hefei, China

## Abstract

Understanding whole-brain circuitry development is essential for uncovering the origins of disorders linked to abnormal neural wiring, yet progress has been limited by the lack of tools for accurate, comprehensive analysis. Brain development is highly dynamic and region-specific, with pronounced inter-individual variability during early stages, and no existing method achieves whole-brain profiling at arbitrary stages. The rapidly changing morphology of developing axons further complicates segmentation. Here, we present D-LMBmapX, a deep learning pipeline for automated whole-brain circuitry profiling across postnatal development. D-LMBmapX constructs sample-inferred atlases from flanking anchor stages, enabling accurate registration at any time point. By developing a foundation model for generalised axon and soma segmentation, it supports quantitative mesoscale developmental connectomics, demonstrated through spatial-temporal profiling of catecholaminergic projections. D-LMBmapX also enables robust cross-modality and cross-dimensional registration, including precise alignment of single 2D slices to 3D references, and offers generalizable strategies for 5D analysis of diverse biomedical spatial datasets.

## INTRODUCTION

In-depth knowledge of neuronal connections is fundamental to understanding how the brain is functionally organised. In the past decades, great success has been achieved in mapping mammalian neural circuits, particularly in mesoscale connectomics^1–5^. Related resources, including high-resolution reference brain atlases, have rapidly expanded ^6–10^. The integration of tissue-clearing techniques with light-sheet fluorescence microscopy (LSFM) has opened up remarkable opportunities for high-throughput, 3D mesoscale connectivity mapping of the entire brain^11–16^. As the proliferation of these large-scale datasets created a critical demand for corresponding analysis tools, we developed D-LMBmap, which employs deep-learning algorithms for whole-brain circuitry profiling, addressing automated axon segmentation, brain region segmentation and whole-brain 3D registration in a single workflow^17^.

Crucially, mapping neuronal connections during development reveals how adult networks are established and provides essential anatomical context for disorders that arise from abnormal wiring^18,19^. Recent whole-brain connectomic analysis of adult and larval Drosophila brains revealed that around 0.5% of neurons have developmental wiring defects, leading to the hypothesis that such miswiring might contribute to individuality and susceptibility to mental health disorders^20^. However, mapping connectivity in developing mammalian brains presents distinct challenges compared to adult brains. In adults, a single time point can serve as a stable reference for circuit architecture. In contrast, developmental connectomics must capture rapid and continuous changes throughout maturation, requiring spatiotemporal mapping across multiple developmental stages and resulting in a workload increase of more than an order of magnitude.

In postnatal mouse brains, structural stabilisation is far from reached before P28^21^. During this period, axonal fibres exhibit diverse and stage-specific morphologies, often differing significantly from those seen in the adult brain^22^. Brain development is highly dynamic and region-specific in the early postnatal period, with each system maturing on its own timeline. For example, the somatosensory cortex receives thalamocortical afferents around P0, and by P3–P4, the critical window for barrel plasticity is largely closed, with barrel hollows and septa clearly defined by P7. In the auditory system, cochlear and brainstem maturation enables the onset of hearing around P10–P12. In the visual system, the eyes open around P12–P14, with functional visual responses beginning immediately thereafter^23^. Other circuits, such as hypothalamic systems, also undergo rapid morphological and synaptic changes during the first two postnatal weeks^24^. To accurately capture such circuit dynamics, both whole-brain registration and axon segmentation must be performed at high temporal resolution—ideally at arbitrary developmental days.

Adding to this complexity is the pronounced inter-individual variability during development. Brain size, structural organisation, and neuronal morphology vary considerably more in developing brains than in adult brains. This variability is especially evident during early postnatal stages, when rapid growth and small differences in developmental timing can produce noticeable anatomical differences, even among littermates. Environmental factors— such as nutrition, maternal care, and minor genetic variation—further amplify these differences.

Current approaches would require repeated atlas construction, whole-brain registration, and axon segmentation on a near-daily basis, involving extensive manual input. This makes the workload impractical and may still result in reduced accuracy due to developmental variability. Moreover, although 3D whole-brain imaging is increasingly available, much circuitry profiling still relies on two-dimensional data, including serial-section methods^10,25–28^, community resources such as the Allen Brain Atlas^29^, and conventional histology. Unlocking the full value of these datasets requires registering 2D information into a 3D framework to enable integration and comparison across modalities.

To address these challenges, we developed D-LMBmapX, a deep learning pipeline for automated whole-brain circuitry profiling at any postnatal stage, offering accurate, efficient analysis through generalised modules adapted to individual samples. D-LMBmapX introduces dynamic 4D and 5D capabilities by incorporating developmental time and multiplexed signal integration. We validated our methods by profiling whole-brain tyrosine hydroxylase (TH)-positive projections and revealed the region-specific and dynamic pattern of TH+ innervations throughout postnatal development. We also achieved robust and automated cross-modality and cross-dimensional registration, demonstrated by aligning genetically defined neuronal populations from the Allen Institute’s developmental ISH datasets^29^ with our 4D TH+ soma maps, refining neuron subtype classification using multiple markers within a unified 5D anatomical-molecular framework. Beyond mouse brains, D-LMBmapX also provides a generalizable strategy for cross-age, cross-modality, and cross-dimensional multiplexed registration across diverse biomedical datasets.

## RESULTS

### A Comprehensive Framework for Developmental Neuronal Circuits Quantification

D-LMBmapX comprises interconnected modules and begins with data input and culminates in automated axon/soma quantification with visualisation in a 3D sample-infered atlas at any postnatal stage (5D, Figure 1). Each core component is powered by advanced deep neural networks (DNNs), and can operate as a stand-alone module or in combination to enable automated 5D whole-brain circuitry profiling, supporting reliable comparison of datasets across platforms.

**Figure 1.**
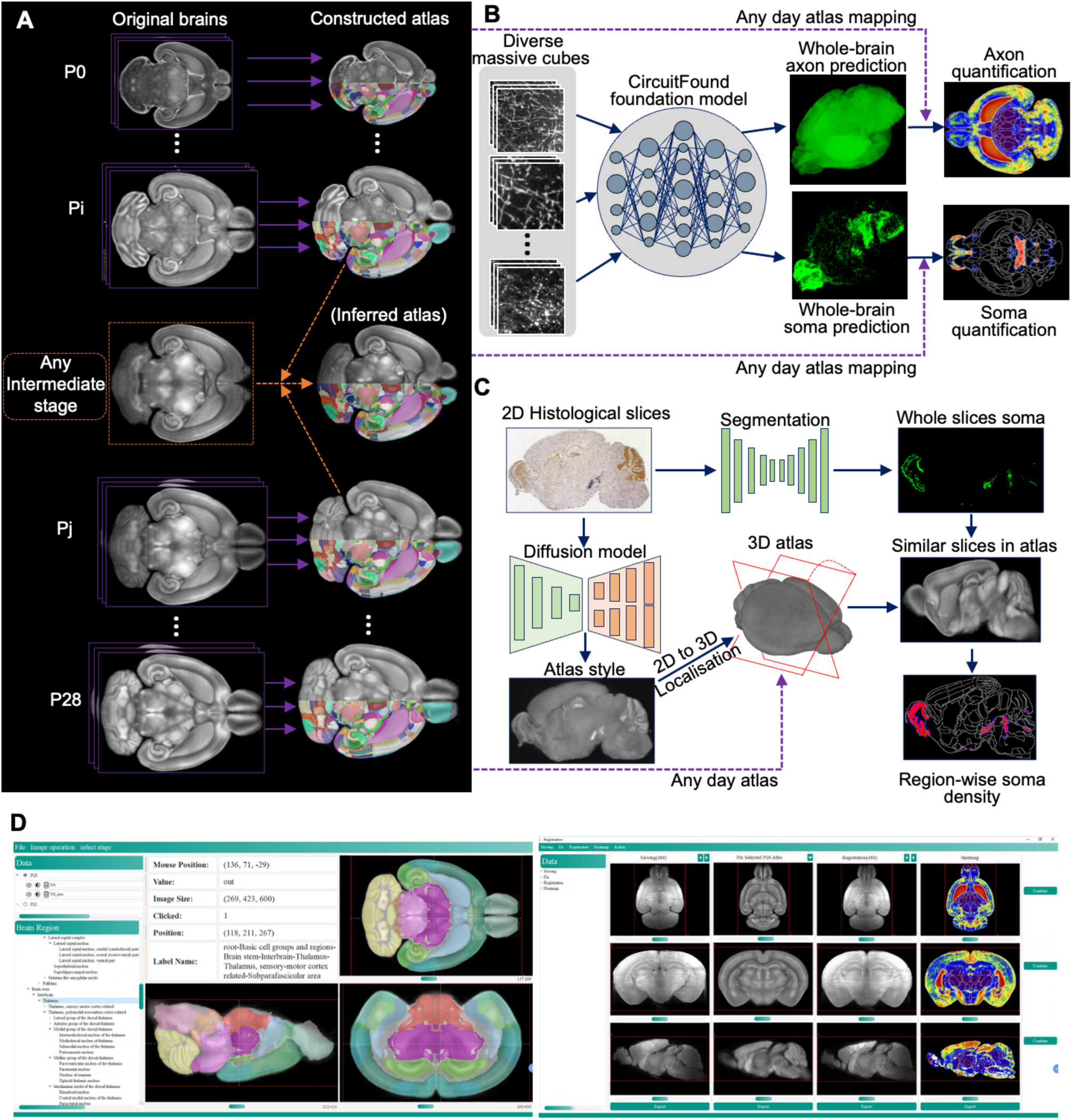
Overview of D-LMBmapX. A. The pipeline for 3D registration of any postnatal day using cross-stage registration. 3D brain atlases were constructed based on LSFM brain samples at postnatal stages P0, P4, P7, P10, P14, P21, and P28. Given an intermediate stage (between Pi and Pj), brain structures can be adaptively quantified through sample-inferred atlas generation and registration with its two flanking atlases. B. The pipeline for quantitative analysis of axons and somata across any developmental stages. Following the extraction of extensive 3D image cubes from whole-brain data, a neuron foundation model is designed to segment axons and somata. C. The pipeline for cross-modality and cross-dimension whole-brain profiling. An example 2D histological slice from the Allen Brain Atlas is used to illustrate the pipeline. The slice undergoes style transfer to match the LSFM modality, followed by automatic localisation and registration of the corresponding section from a digitally sliced 3D atlas. Soma segmentation is then performed to enable region-wise quantification of soma density. D. The software interface of the D-LMBmapX for neural circuitry profiling and atlas visualisation of developing mouse brains.

Using a novel registration approach, we first constructed high-resolution average template brains at seven anchor postnatal stages (P0, P4, P7, P10, P14, P21, and P28). Hierarchical brain structures were then delineated by sequentially registering the Allen Mouse Brain Common Coordinate Framework (CCFv3)^7^ to these templates. To address stages not covered by the seven anchor atlases, we developed a sample-inferred atlas construction strategy. This approach adaptively delineates brain regions by inferring an atlas from flanking anchor stages using even a single sample, enabling accurate whole-brain registration at any postnatal time point (Figure 1A). To achieve whole-brain axon and soma profiling throughout development, we developed a generalised foundation model CircuitFound for automated segmentation, capable of accurately segmenting axons and somata despite morphological differences across stages. Coupled with the whole-brain registration model, this allows D-LMBmapX to accept input from any postnatal brain and output region-specific quantification and 3D visualisation of axon and soma distributions, registered to the appropriate developmental atlas (Figure 1B). This cross-dimensional and cross-modality module supports end-to-end quantitative analysis of 2D histological sections via similarity-driven 2D-to-3D spatial alignment and registration—estimating each section’s pose, refining with deformable alignment to correct tissue distortion, and mapping quantitative signals into atlas coordinates for region-wise analysis (Figure 1C).

D-LMBmapX integrates these components into a user-friendly software platform that is open-source, supports customisation through high-level application programming interfaces (APIs), and includes a graphical interface for selecting and executing pretrained deep learning models (Figure 1D).

### Multi-Constraint Non-Rigid Pipeline for 4D Brain Atlas Construction

To achieve whole-brain registration across any postnatal stage, we first constructed high-resolution 3D brain atlases for seven anchor stages. We collected LSFM brain images from P0 to P28, and used at least 30 half-brains per stage to construct each average template. To minimise variability from sample preparation and imaging, we applied a series of image pre-processing strategies, including N4 bias field correction^30^, rescaling intensity, and 3D adaptive histogram equalisation^31^. For each stage, we created an initial reference template by aligning all brains to a common centre and computing the voxel-wise average (Figure 2A). While we incorporated the conventional atlas-building pipeline^32,33^—which includes rigid, affine, and B-spline registration—we further refined the templates using a custom 2.5D non-rigid registration approach. It uses a U-Net-based neural network to perform 2D registrations in both coronal and horizontal planes. We then assembled the 3D registration results from each view and computed a 3D consistency loss to guide iterative refinement. This process allows each brain to update the template within a shared deformation space, resulting in a high-resolution, fine-grained reference template (Figure 2B).

**Figure 2.**
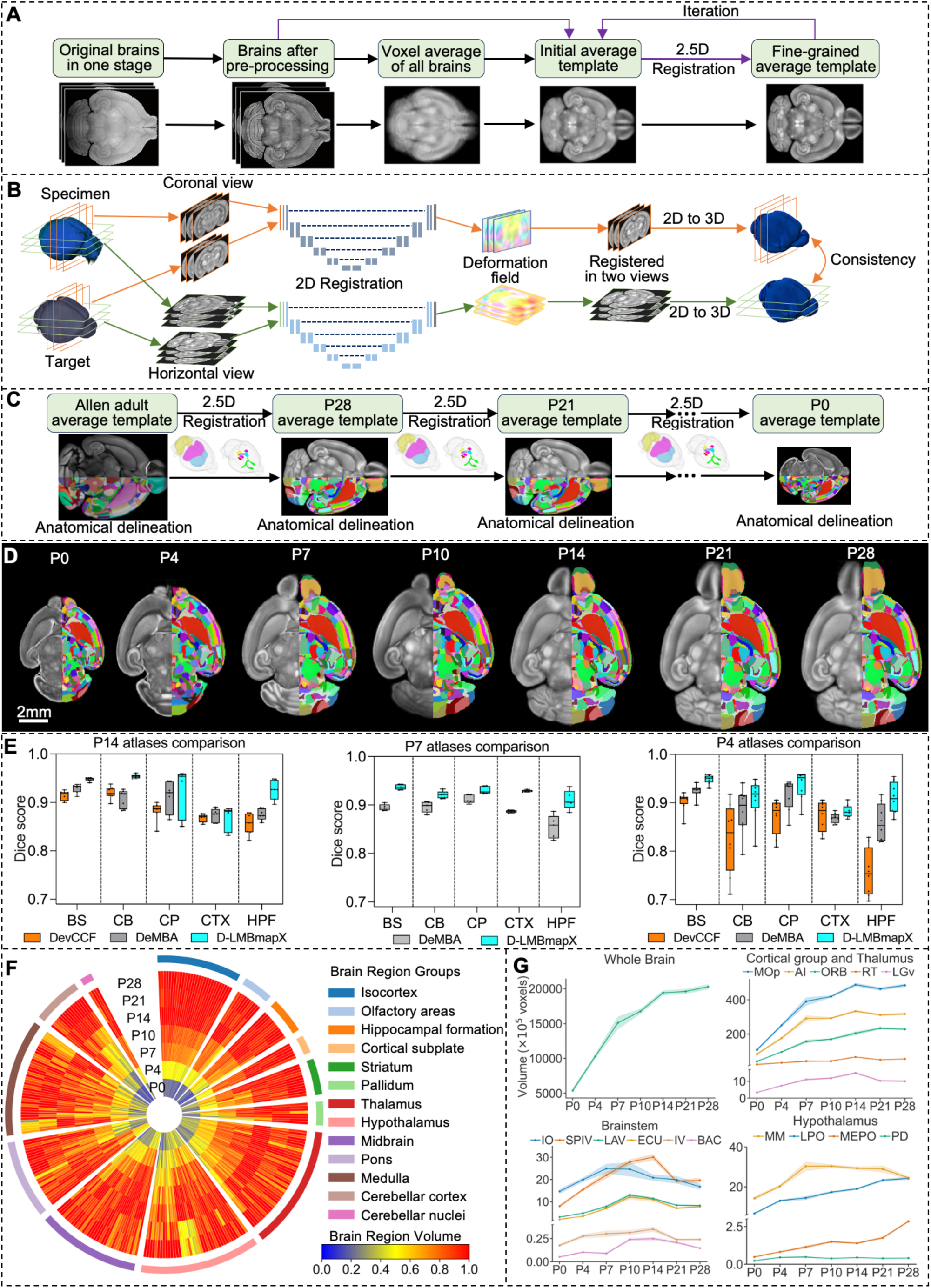
Construction of 3D mouse brain atlas across seven postnatal stages. A. The pipeline for the construction of 3D averaged brain in each developmental stage. Original brain samples from each stage are first pre-processed and voxel-wise averaged. The individual brain was registered to the voxel-wise averaged brain with rigid, affine, and deformable transformations to generate the initial reference template. Finally, a fine-grained average template was generated through iterative registration of pre-processed brains to the initial reference template using the developed 2.5D registration method. B. Multiview 2.5D registration framework. The coronal and horizontal views of the input 3D brain sample are first registered to the corresponding 2D image slices of the reference brain. Deformation field obtained from two views are used for registration. 3D Registered results from either views are used for the computation of the 3D consistency loss. C. The pipeline of brain atlas construction based on fine-grained average templates. The P28 atlas was delineated from Allen CCFv3, and subsequent stages (P21–P0) were sequentially derived, with each delineation performed using 2.5D registration with multiple brain region constraints. D. Visualisation of the 3D developmental brain atlases across seven postnatal stages in horizontal view. For each atlas, the left side is shown with the average brain template and the right half with anatomical delineations. E. Evaluation and comparison of three developmental mouse brain atlases, DevCCF, DeMBA and D-LMBmapX. Performance was assessed by calculating region-wise Dice scores for major brain areas (BS, CB, CP, CTX, HPF) after registering LSFM sample brains to each atlas at P14 (n=7), P7 (n=5), and P4 (n=8). Box plot: centre line, median; box limits, upper and lower quartiles; whiskers, 1.5× interquartile range. F. Radial plot illustrating the temporal progression in anatomical volume changes of 276 profiled brain regions across seven postnatal stages. Individual brain regions are grouped according to their boarder anatomical categories (Table. S5). Colour code indicates scaled volume changes across developmental stages for each brain region. G. Volume changes of the whole brain and selected brain regions across postnatal development. Top left, whole brain; top right, selected brain regions from cortical and thalamic areas; bottom left, selected brain regions containing dopaminergic neurons; bottom right, selected brain regions from the hypothalamus (n=8 brains in each stage; solid lines and shadows represent mean ± SEM).

With fine-grained brain templates established for each developmental stage, our next challenge was to delineate hundreds of brain regions for each one. Manual annotation at this scale is extremely labour-intensive, so we developed an automated pipeline for atlas mapping and construction using cross-stage and cross-modality registration (Figure 2C). We began by registering the adult mouse brain atlas, Allen CCFv3, to our P28 brain template. This process combined brain style transfer, multi-constraint registration^17^, and our 2.5D non-rigid registration approach. Specifically, we developed diffusion-based brain style transfer to convert the Allen CCFv3 images which are in STPT modality into the LSFM modality. We created multiple brain structure constraints by automatically annotated six major brain regions along with manual annotated eight smaller regions (Figure S1A). The multi-constraint strategy alleviated the alignment shifting and registration error accumulation problem during the propagation of brain region delineation (Figure S1B). Following cross-modality and cross-stage alignment from Allen CCFv3 to the P28 template, the pipeline automatically delineated hundreds of brain regions. We then propagated these annotations to earlier stages through sequential registration—P28 to P21, P21 to P14, and so on—ultimately constructing a complete atlas series across all seven postnatal stages (Figure 2D, Figure S2).

To evaluate the effectiveness of our 4D brain atlas construction pipeline and assess the accuracy of the resulting D-LMBmapX atlas, we conducted a series of comparative analyses against established methods and reference atlases. We first benchmarked our whole-brain registration pipeline against ANTs^34^, a widely used non-rigid registration method which was also adopted in constructing the DevCCF atlas^32^. Region-wise Dice scores were computed across the entire brain following registration to evaluate anatomical alignment accuracy. For a fair comparison, we incorporated the same pre-processing into both pipelines. Our results demonstrate that D-LMBmapX consistently achieves higher Dice scores across multiple brain regions, indicating the superior performance of our 2.5D non-rigid registration approach (Figure S1C). We then assessed the accuracy of atlases constructed using D-LMBmapX by comparing them with two leading developmental brain atlases: DevCCF^32^ and DeMBA^35^. Because these atlases were originally built using different registration strategies, we re-registered independent brain samples to each atlas using the same D-LMBmapX registration pipeline to ensure a fair comparison. Since DevCCF and DeMBA^35^ do not cover all the postnatal stages available in D-LMBmapX, we restricted the evaluation to the stages present in all three atlases. For DeMBA^35^, only the authentic (real-sample) atlases were included in the analysis, excluding their interpolated (synthetic) ones. Region-wise Dice scores were then used to assess anatomical alignment, with D-LMBmapX showing consistently higher anatomical correspondence (Figure 2E, Figure S3).

Together, these results validate the effectiveness of our multi-constraint non-rigid pipeline for whole-brain registration and demonstrate that D-LMBmapX produces a high-accuracy 4D brain atlas capable of capturing developmental dynamics with improved anatomical precision over existing resources.

Next, we characterised developmental volume changes at the whole-brain and regional levels across seven postnatal stages (Figure 2F and G, Table S6). At the whole-brain level, total volume increased steadily, with the fastest growth between P0 and P7 (Figure 2G). At the regional level, 276 regions grouped into 13 major brain areas showed distinct and dynamic developmental trajectories (Figure 2F). The cortical and hippocampal areas, striatum, and cerebellum broadly exhibited monotonic increases, although cortical subregions varied: the agranular insular area (AI) plateaued at P7, the primary motor cortex (MOp) at P14, and the orbital area (ORB) at P21 (Figure 2G). By contrast, thalamic regions peaked around P14, while other major brain groups displayed more heterogeneous, stage-specific patterns of change (Figure 2F).

These findings underscore that brain maturation is both nonlinear and regionally asynchronous, with different brain areas undergoing distinct developmental trajectories and reaching critical milestones at varying postnatal timepoints—some of which may fall outside the seven anchor stages profiled here. This biological complexity necessitates whole-brain registration at arbitrary postnatal days to ensure anatomical accuracy. However, the interpolation-based strategy used in DeMBA^35^, which generates intermediate-stage atlases by assuming smooth and uniform brain development between flanking timepoints, does not account for these region-specific dynamics. As a result, such interpolated atlases may introduce anatomical inaccuracies, particularly during periods of rapid or heterogeneous development.

### Sample-inferred Whole-brain Registration at Any Time Point

To enable accurate whole-brain registration at any postnatal stage, we developed a pipeline that constructs a sample-inferred atlas by integrating the brain sample under study with atlases from its adjacent developmental stages (Figure 3A). To demonstrate the pipeline, we use a P10 brain as an example, constructing a sample-inferred P10 atlas from its flanking stages (P7 and P14). First, the P7 and P14 atlases were registered to the P10 brain image, generating two deformed atlases in a P10-like space while preserving their original anatomical labels. These P10-like atlases were then voxel-wise averaged to form an initial reference template that integrates anatomical features from both flanking stages. Following the atlas construction pipeline illustrated in Figure 2A, the two P10-like atlases were iteratively registered to the initial average template, producing a refined sample-inferred P10 atlas with improved anatomical consistency (Figure S4A). Next, to delineate the brain sample, the original P10 brain was registered to this sample-inferred atlas, and all anatomical labels were transferred. To further enhance registration reliability and general applicability, we designed a mutual learning–based registration framework that automatically identified reliable anatomical constraints. Specifically, brain regions showing strong spatial correspondence between the two sources (P10-like P7 and P10-like P14 atlases) were automatically selected and used as constraints to guide registration from the original P10 brain to the sample-inferred atlas. Here, two 2.5D registration models were mutually trained based on constraints from the intersection and union of selected brain regions, respectively (Figure S4B).

**Figure 3.**
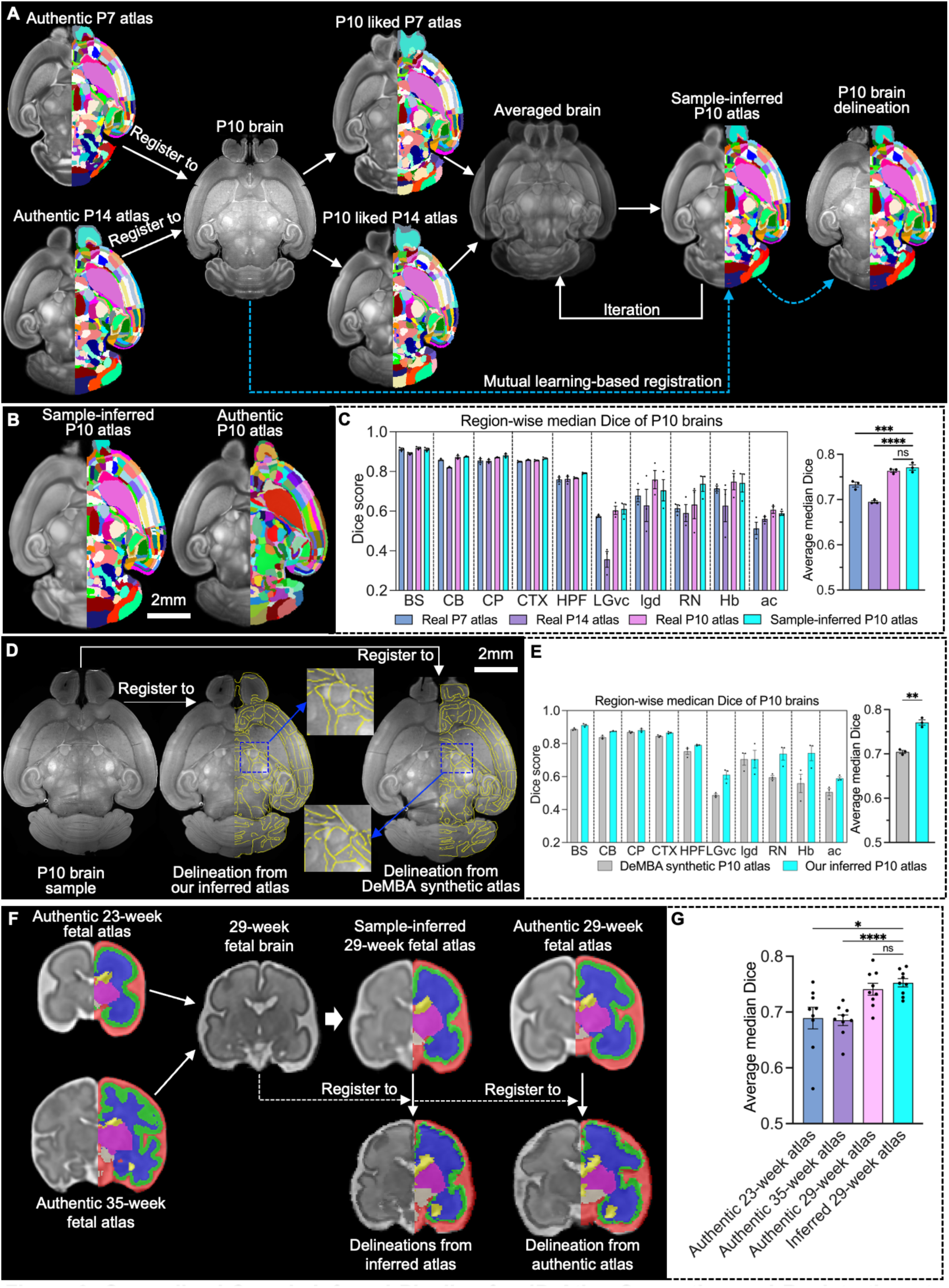
Generalised Sample-Inferred Pipeline for 4D Atlas Generation and Registration at Arbitrary Developmental Stages. A. Pipeline for constructing a sample-inferred 3D brain atlas at any developmental stage using flanking atlases — illustrated with a P10 mouse brain as an example. When a P10 brain sample needs to be registered but only P7 and P14 atlases are available, D-LMBmapX constructs a sample-inferred P10 atlas. The P7 and P14 atlases are non-linearly registered to the P10 brain, averaged, and iteratively refined into a consistent inferred template. The P10 brain is then registered to this atlas, enabling full-brain anatomical delineation through label transfer. B. Representative horizontal view of the sample-inferred P10 atlas and the authentic P10 atlas constructed by D-LMBmapX. The sample-inferred P10 atlas was constructed using a single P10 brain and two flanking atlases (P7 and P14), following the pipeline described above. The authentic P10 atlas was constructed from 30 P10 brain samples following the standard template construction pipeline shown in Figure 2. C. Quantitative comparison of P10 registration to sample-inferred, authentic, and flanking-stage atlases. Left, region-wise Dice score for major brain regions (BS, CB, CP, CTX, HPF) and small brain regions (LGvc, lgd, RN, Hb, ac). Right, average median Dice score after registering the sample brain to each atlas (calculated by ordinary one-way ANOVA with Dunnett’s multiple comparison, n=3 brains. ns indicates non-significant, ***P<0.001, ***P<0.0001). Measure of centre, mean; error bars, mean±SEM. D. Representative horizontal view of a P10 sample brain registered to the DeMBA interpolated P10 atlas and the D-LMBmapX sample-inferred P10, respectively, with orange lines marking anatomical region boundaries in each atlas. E. Quantitative comparison of P10 sample brains registration to the DeMBA interpolated P10 atlas and the D-LMBmapX sample-inferred P10. Left, region-wise Dice score for major brain regions (BS, CB, CP, CTX, HPF) and small brain regions (LGvc, lgd, RN, Hb, ac). Right, average median Dice score after registering sample brains to each atlas (calculated by Two-tailed paired t-test, n=3 brains. **P<0.01). Measure of centre, mean; error bars, mean±SEM. F. Whole-brain registration of a 29-week human fetal brain to the sample-inferred atlas and authentic atlas. Construction of the 29-week human fetal brain sample-inferred atlas was based on the authentic flanking atlases at 23-week and 35-week. G. Quantitative comparison of a 29-week human fetal brain registration to the sample-inferred 29-week atlas, authentic 29-week atlas, and two flanking 23-week and 35-week atlases. Average median Dice score after registering sample brains to each atlas (calculated by one-way ANOVA with Dunnett’s multiple comparisons, n=9 brains, *ns* indicates non-significant, \**P*<0.05 ****P<0.0001). Measure of centre, mean; error bars, mean±SEM.

This strategy leveraged reliably aligned brain regions as spatial constraints to boost registration performance across the whole brain (Figure S4C). Importantly, all brain-region constraints—including both major and small regions—were obtained automatically, without any manual annotation. To assess similarity between the sample-inferred and authentic P10 atlases (Figure 3B), P10 brain samples were registered to each, as well as to the flanking-stage atlases (P7 and P14). Dice score analysis revealed no significant differences in registration accuracy between the sample-inferred and authentic P10 atlases for either major or small brain regions, whereas registration to P7 or P14 yielded substantially lower accuracy (Figure 3C). Comparable results were obtained when P7 was used as the inferred stage (Figure S5A & B). Next, we compared the registration performance of a P10 brain sample aligned to the interpolated P10 atlas from DeMBA versus our sample-inferred P10 atlas. Dice score analysis showed that registration to our atlas yielded superior accuracy both globally and across individual brain regions (Figure 3D & E).

Collectively, these results demonstrate that our sample-inferred whole-brain registration pipeline enables accurate registration at any postnatal day. The resulting sample-inferred atlases achieve registration accuracy comparable to authentic atlases and substantially exceed that of interpolated alternatives.

### Generalised Pipeline for Sample-Inferred 4D Atlas Generation and Registration

Our sample-inferred registration pipeline generalises beyond the developing mouse brain and supports 4D analysis of human fetal and infant datasets. For both a 29-week fetal brain and a 7-month infant brain, we generated a sample-inferred atlas by integrating the sample itself with its flanking-stage atlases (e.g., 23- and 35-week, or 4- and 10-month). Each brain was then registered to its own sample-inferred atlas, the corresponding authentic atlas, and the flanking-stage atlases. Dice score analysis showed that registration to the sample-inferred atlas achieved accuracy comparable to the authentic atlas and significantly exceeded that of flanking-stage registration (Figure 3F & Figure S6).

### 5D Whole-brain Circuitry Profiling by a Generalised Foundation Model

Whole-brain circuitry profiling requires effective segmentation of neuronal structures (e.g., axon and soma) by deep learning models trained on abundant, well-annotated samples. To eliminate the labour-intensive manual annotation of axons, previously we developed automated annotation and data augmentation strategies^17^. However, neuronal structures, especially axonal projections, undergo significant morphological changes throughout development. This morphological diversity poses a major challenge for robust circuitry profiling across stages. Segmentation modules trained on a single stage often fail to generalise, and training new models for every stage is too costly in time and resources. To address this challenge, we developed CircuitFound, a generalised foundation model integrated into D-LMBmapX for axon and soma segmentation that adapts effectively to diverse neuronal morphologies and brain images (Figure 4).

**Figure 4.**
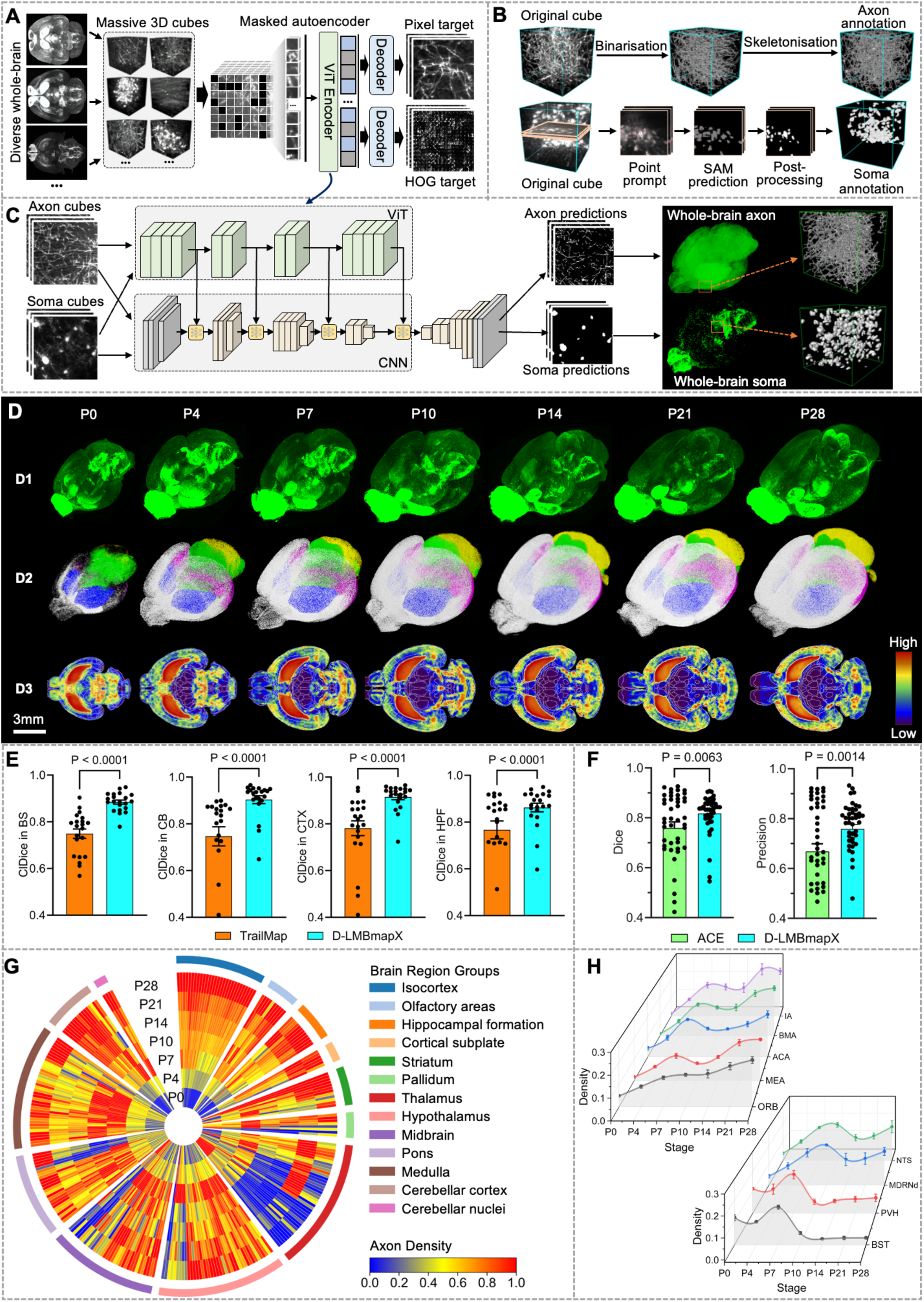
CircuitFound, a generalised foundation model for 4D whole-brain circuitry profiling. A. The self-supervised pre-training framework designed in CircuitFound. Over 100,000 3D cubes were randomly cropped from LSFM brain samples. A customised dual-decoder 3D masked autoencoder (MAE) was trained from scratch in a self-supervised manner. In addition to reconstructing voxel-level cubes, the network also reconstructed cubes after computing histograms of oriented gradients (HOG), enabling improved capture of long-range, tree-like axon morphologies. B. Automated and semi-automated annotation of axon and soma in selected sample cubes. Axon annotation was performed using the binarisation and skeletonisation pipeline embedded in D-LMBmap. Soma annotation was carried out with a newly developed pipeline combining manual 2D point prompting, prediction with the Segment Anything Model (SAM), and post-processing. C. Task-specific fine-tuning network in CircuitFound. With a small number of task-specific cubes containing automated annotations (axons or somata), features were extracted using the ViT encoder from the pre-trained CircuitFound model. A multi-level cascading CNN module was then introduced in parallel with the ViT encoder, enabling faster convergence and robust performance with only a few fine-tuning samples. D. Whole-brain axon segmentation and projection mapping of anti-TH stained brains in postnatal stages from P0 to P28. Top row, representative LSFM brains across seven stages in the anti-TH stained channel. Middle row, representative results of whole-brain segmentation of TH+ axon across seven stages, where axons located in different major brain regions are visualised in distinct colours. (white: CTX and OLF, blue: CP, ACB, and OT, green: BS, yellow: CB, pink: HPF). Bottom row, heatmaps of averaged TH+ axon distribution in horizontal view from P0 to P28 (n=3 for each stage). E. Quantitative comparison of axon segmentation effectiveness between TrailMap and D-LMBmapX by the ClDice metric. A total of 46 annotated cubes (150 × 150 × 150) were used for training both models. For evaluation, 84 cubes were sampled from four major brain regions (BS, CB, HPF, and CTX) across 21 anti-TH–stained brains, with three brains assessed at each developmental stage (P0, P4, P7, P10, P14, P21, P28). Two-tailed paired *t*-test. Error bars represent mean ± SEM. F. Quantitative comparison of soma segmentation effectiveness between ACE and D-LMBmapX under the evaluation metric of Dice and Precision. A total of 36 annotated cubes (150 × 150 × 150) were used for training both models. For evaluation, 44 cubes were sampled from 21 anti-TH–stained brains across seven developmental stages. Two-tailed paired t-test. Error bars represent mean ± SEM. G. Radial plot illustrating the temporal progression in TH+ axon density during postnatal development across 276 profiled brain regions. Individual brain regions are grouped according to their boarder anatomical categories (Table. S6). Colour code indicates scaled axonal density across developmental stages for each brain region. H. Waterfall plot showing the TH+ axon density changes during postnatal development for selected brain regions. Top, selected brain regions where TH+ axon density is significantly increased in the other six postnatal stages compared to P0. Bottom, selected brain regions where TH+ axon density is increased in other six postnatal stages compared to P7 (One-way ANOVA followed by Bonferroni correction. P value in Table S8), n=3 brains for each stage. Error bars, mean±SEM.

The construction of CircuitFound contained three steps: self-supervised pre-training, automated sample annotation, and fine-tuning. More than 100,000 3D cubes randomly cropped from anti-TH stained LSFM brain samples across seven postnatal stages are used for self-supervised pre-training by a 3D masked autoencoder (MAE). A Vision Transformer (ViT)^36^ encoder embeds the unmasked patches, and unlike conventional MAEs that rely solely on pixel-level reconstruction^37^, we introduced a second decoder branch to reconstruct 3D Histograms of Oriented Gradients (HOG)^38^ (Figure 4A). This dual-decoder design enables the model to capture both general image features and gradient-based structural cues, allowing it to better represent the long-range, tree-like morphology of axons critical for accurate segmentation. Ablation study results support the effectiveness of our strategies in pre-training (Figure S7A).

Many neural systems, such as the dopaminergic system, have somata distributed across multiple brain regions rather than confined to a single nucleus. To accurately reconstruct circuitry, segmentation must therefore distinguish somatic signals from axonal projections, as conflating the two can lead to misinterpretation of connectivity. Thus, CircuitFound is designed to characterise both axons and somata, and after pre-training, both are annotated automatically. We used the previously reported method in D-LMBmap for axon annotation, and developed a new pipeline for 3D soma annotation based on the segment anything model (SAM)^39^. In this pipeline, the only manual input is a point prompt for each soma in a few 2D slices; SAM then automatically segments the somata from these prompts. To further separate adhered somata, we incorporated a watershed algorithm based on distance maps (Figure 4B). This lightweight strategy enables efficient and extensible annotation for diverse axon and soma segmentation tasks. Lastly, we fine-tuned the pre-trained foundation model with annotated cubes. While directly fine-tuning the ViT encoder is inefficient and data-hungry due to its large number of parameters, we addressed this by introducing a parallel convolutional neural network (CNN). At each level, the ViT encoder output is fused with the corresponding CNN output, enabling faster convergence and improved accuracy (Figure 4C). By paralleling ViT and CNN, our framework achieves robust whole-brain prediction of axons and somata with only a small number of annotated samples.

Equipped with the CircuitFound module, D-LMBmapX delivers robust performance in generalisation for axon and soma segmentation. To demonstrate its effectiveness in whole-brain circuitry profiling across brain regions and developmental stages, we applied it to characterise catecholaminergic circuits over seven postnatal stages. Brains from P0, P4, P7, P10, P14, P21, and P28 mice were processed with whole-mount immunohistochemistry, iDISCO tissue clearing, and LSFM imaging. We used tyrosine hydroxylase (TH) immunostaining to label dopaminergic and noradrenergic neurons and their axonal projections. D-LMBmapX successfully segmented these labelled structures, registered each brain to its corresponding developmental stage atlas, and computed the density map of TH+ axons and somata for each stage (Figure 4D, Figure S8, Video S1-3). In comparison with traditional deep models such as TrailMap^12^, D-LMBmap^40^ and ACE^41^, D-LMBmapX doesn’t have to do sample training for the individual stage. Moreover, D-LMBmapX achieves superior performance, especially when using a smaller number of annotated samples (Figure S7B). For axon segmentation, D-LMBmapX consistently outperformed TrailMap in ClDice, ClPrecision, ClRecall and Dice scores across different brain regions and whole brain on average at any developmental stages (Figure 4E and Figure S7C). For soma segmentation, D-LMBmapX significantly outperformed ACE, which is the most advanced pipeline for mapping neuronal cell bodies, in Dice and Precision scores (Figure 4F).

Using D-LMBmapX, we quantified TH⁺ axon densities across more than 260 brain regions spanning 13 major brain areas and generated a 4D atlas of postnatal axonal innervation (Figure 4G, Table S6, Video S2&3). This analysis revealed highly heterogeneous, region-specific patterns of TH⁺ innervation. In the isocortex and olfactory areas, axonal density broadly increased over time, whereas other regions displayed pronounced spatiotemporal dynamics (Figure 4G–H). These included sharp increases at specific developmental stages (e.g., NTS), transient peaks (e.g., PVH and BST), stable levels from birth, or even a progressive decline (Figure 4H, Figure S9). To our knowledge, this is the first 4D whole-brain map of TH⁺ axonal innervation. Notably, decreases in axonal density could not be fully explained by regional volume changes, suggesting a contribution from axonal pruning (Figure S9).

### Generalized Cross-dimensional and Cross-modality Registration

Although 3D whole-brain imaging is increasingly adopted, much of the available circuitry profiling data remains two-dimensional. This includes advanced imaging techniques that acquire data in serial sections, major community resources such as the Allen Brain Atlas, and the widespread use of 2D histology in many neuroscience laboratories. To ensure the broader neuroscience community can fully benefit from diverse connectivity mapping resources and enable cross-platform data comparison, we developed cross-dimensional and cross-modality registration within D-LMBmapX.

The cross-dimensional and cross-modality registration pipeline includes slice pre-processing, cross-modality style transfer, spatial alignment to identify the best-matching section in a 3D reference atlas, and final 2D-to-3D registration (Figure 5A). Previously, we developed the first brain-specific style transfer method based on CycleGAN, which learns an unsupervised mapping between imaging modalities^17^. However, this approach requires retraining the model from scratch whenever a new modality is introduced. To have more generalized and consistent cross-modality style transfer, we developed a new solution based on the diffusion model, which can transfer source brain image modality to any given target style (Figure S10A). During training, only target brain images are used. Each image is gradually degraded by adding noise until it becomes indistinguishable from random noise. The model then learns to reconstruct the original target image through a denoising process.

**Figure 5.**
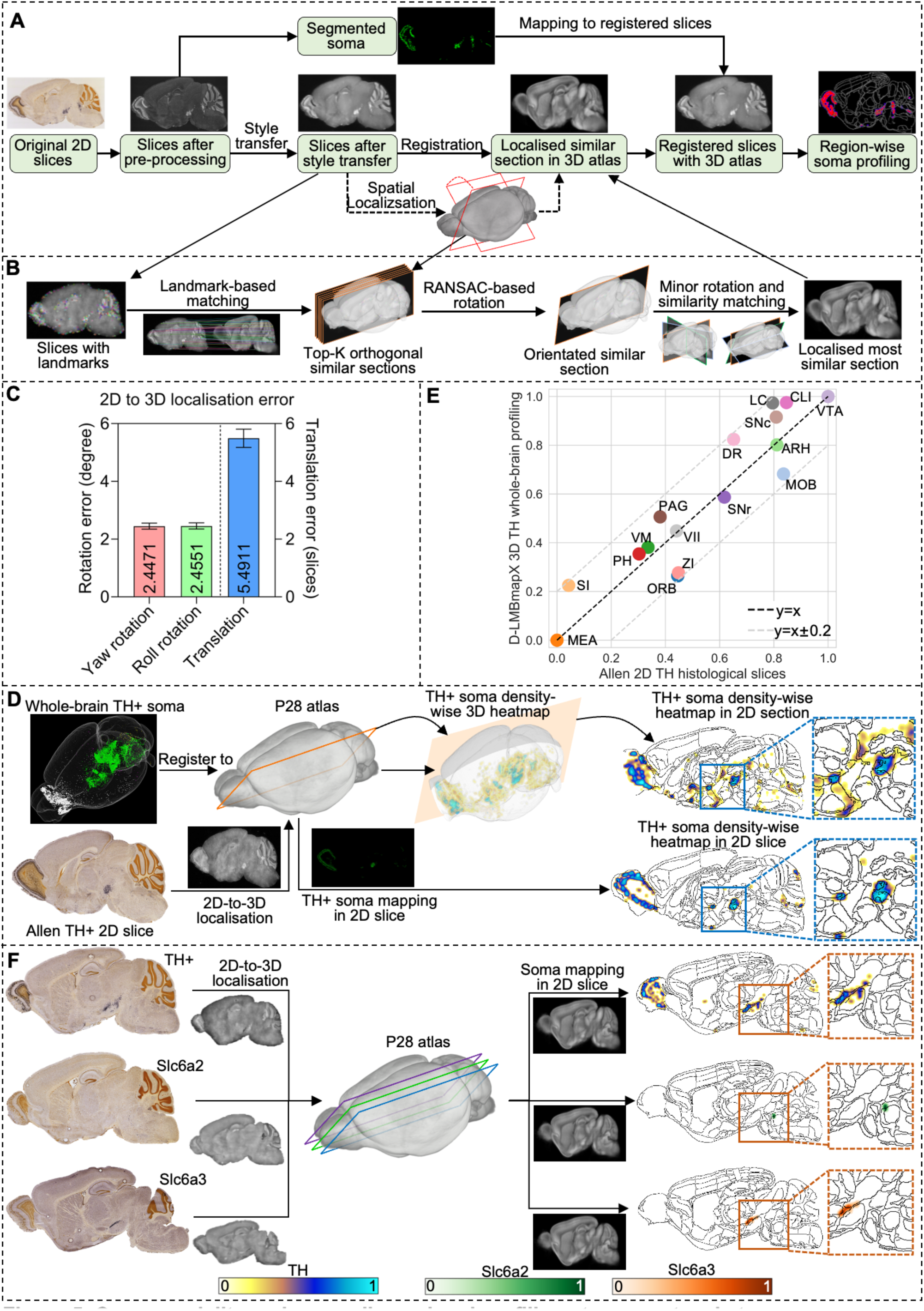
Cross-modality and cross-dimensional profiling at any postanal stages. A. Automated pipeline for comprehensive 2D-to-3D brain mapping and region-wise quantification. It includes slice pre-processing, cross-modality style transfer, and spatial alignment to identify the best-matching sections in a 3D reference atlas. This is followed by 2D-to-3D registration, automated soma/axon segmentation, and region-wise quantitative analysis. B. Design of the automated 2D-to-3D spatial alignment module. Corner-based landmarks are extracted from style-transferred 2D histological slices and from all orthogonal sections (e.g., sagittal) of the 3D reference atlas. Top-K candidate sections are selected based on matched landmarks (K=40 in our experiment). Initial orientation is estimated using RANSAC to align 2D slice landmarks with those from top-K sections. D-LMBmapX then refines localisation by applying small yaw and roll adjustments, selecting the best-matching section using a Mutual Information (MI) metric. C. Evaluation of 2D-to-3D spatial localization accuracy. The Allen 3D CCFv3 atlas (STPT modality) was first registered to the D-LMBmapX P28 3D atlas (LSFM modality) to establish ground truth alignment. A total of 169 2D slices were randomly selected from the Allen CCFv3 STPT atlas and subjected to arbitrary yaw (rotation in Z-axis), roll (rotation in Y-axis), and translation (linear shifts along x, y, or z). Each slice was then spatially aligned to the 3D LSFM atlas using the D-LMBmapX pipeline. Localisation accuracy was assessed by comparing yaw, roll, and translation errors against the ground truth. D. Comparison of TH⁺ somata density in corresponding sagittal panels from two sources: P28 Allen 2D ISH slices (top) and matched sections extracted from a P28 3D anti-TH–stained LSFM brain (bottom) profiled by D-LMBmapX. The 3D whole-brains stained with TH^+^ somata were registered to the P28 LSFM atlas to profile the somata density in 3D. The P28 Allen 2D TH^+^ slices were localised to the 3D P28 atlas in matching similar sections, combining with somata segmentation to achieve the TH^+^ somata profiling in 2D slices. E. Evaluation of consistency in normalised region-wise TH⁺ soma density between Allen 2D ISH slices and 3D LSFM whole-brain data profiled with D-LMBmapX at P28. F. Examples of registering a single 2D slice into 3D brain atlas. Selected P28 Allen 2D ISH slices for three gene markers (*Th*, *Slc6a2*, and *Slc6a3*), each containing the ventral tegmental area (VTA) and locus coeruleus (LC), are profiled. Left: original Allen ISH slices for *Th*, *Slc6a2*, and *Slc6a3*. Middle: corresponding sections localised, registered, and mapped within the P28 LSFM 3D atlas. Right: heatmaps of region-wise soma density for *Th*, *Slc6a2*, and *Slc6a3*. Soma segmentation, 2D-to-3D registration, and region-wise density profiling were performed using D-LMBmapX. Heatmap colours indicate normalized somata density per sample, with colour scales shown in the corresponding bars for each gene.

To improve stability and accuracy, we incorporated two modules: (i) brain edge constraints, which ensure structural boundaries in the reconstructed image remain consistent with the input, and (ii) a denoising U-Net, which segments major brain regions (BS, CB, CTX, HPF, CP, OLF) to preserve regional boundaries. In the prediction step, source brains from other modalities are transformed into the target style using the trained model, with structural edges from the source guiding reconstruction to maintain anatomical detail. This diffusion-based approach offers two major advantages: it preserves edges and fine details across modalities, and it requires training only once for a given target modality, independent of source modalities. Benchmark comparisons show that D-LMBmapX achieves superior performance in brain style transfer, including transformations from LSFM to STPT (Allen CCFv3) and from Allen histological 2D slices to our developmental LSFM brain atlas (Figure S10B & C).

After preprocessing and style transfer to match the target 3D brain modality, each 2D slice was automatically localised and registered within the 3D atlas using our newly developed modules (Figure 5B). Landmarks from both the 2D slice and each orthogonal section of the 3D atlas (e.g., sagittal view) were automatically detected using the Harris corner detector^42^ and described with Scale-Invariant Feature Transform (SIFT)^43^ features. The top-K most similar orthogonal sections were identified based on the number of matched landmarks, and the Random Sample Consensus (RANSAC) algorithm^44^ was applied to estimate the optimal initial rotation aligning the 2D slice to these candidate sections. To refine alignment, D-LMBmapX further performed fine-grained searches around the estimated orientation by varying yaw and roll angles, selecting the best-matched section according to mutual information similarity. To assess localisation accuracy, we randomly perturbed 2D slices with controlled rotations and translations and then aligned them to the 3D atlas using D-LMBmapX. The resulting low errors across yaw, roll, and translation confirmed the robustness and precision of our pipeline for reliable 2D-to-3D registration (Figure 5C).

To demonstrate that D-LMBmapX enables accurate 2D-to-3D whole-brain registration for systematic circuitry profiling, we characterised and registered marker gene expression related to catecholaminergic systems. We first compared Th⁺ soma density between Allen 2D ISH slices and anti-TH–stained 3D LSFM brains at P4, P14, and P28 (Figure 5D, Figure S11). Across 16 brain regions containing dopaminergic or noradrenergic neurons, normalised region-wise TH⁺ soma densities were well aligned between the two datasets (Figure 5E). Next, to further refine cell-type characterisation by integrating 2D molecular signatures with 3D anatomical organisation, we profiled and registered individual Allen 2D ISH slices encompassing both the ventral tegmental area (VTA) and locus coeruleus (LC) for three gene markers—*Th*, *Slc6a2*, and *Slc6a3*—at P4, P14, and P28. Heatmaps of soma density generated by D-LMBmapX accurately localised dopaminergic neurons in the VTA (*Th*⁺, *Slc6a2*⁻, *Slc6a3*⁺) and noradrenergic neurons in the LC (*Th*⁺, *Slc6a2*⁺, *Slc6a3*⁻)^45^ (Figure 5F, Figure S12). Together, these results show that D-LMBmapX can accurately place single 2D histological sections within a 3D atlas, robustly matching them despite differences in imaging position, rotation, or deformation, and even without continuous serial sections. To our knowledge, this represents the first workflow to achieve automated cross-dimensional and cross-modality whole-brain registration, providing a generalised framework for integrating diverse spatial biomedical images.

## DISCUSSION

The rapid rise of mesoscale connectomics, coupled with advances in machine learning, has accelerated the shift toward registering circuitry and multi-omic data to standardised brain atlases. However, compared to the growing wealth of tools and resources available for the adult brain^46–51^, equivalent resources for developing brains remain scarce. Here, we present D-LMBmapX, an end-to-end deep-learning system for 5D mapping of mouse brain circuitry across postnatal development. D-LMBmapX includes automated tools for whole-brain registration and soma/axon segmentation across all postnatal stages, enabling efficient and robust quantitative analysis of developmental connectomics. It further offers generalised strategies for segmentation and registration across temporal, spatial, and modality domains, supporting scalable and versatile applications to a wide range of biomedical datasets.

We developed a custom 2.5D non-rigid registration strategy that enables accurate whole-brain registration between individual samples and 3D references, thereby facilitating efficient construction of anchor-stage atlases. Unlike typical 3D deep learning–based approaches, our 2.5D registration approach achieves accurate, high-resolution 3D registration without incurring heavy computational costs. This strategy is readily scalable to higher-resolution images, enabling broader applicability across diverse datasets. With it embedded in D-LMBmapX, we constructed seven anchor-stage atlases and quantified volumetric changes in 267 brain regions across postnatal development. This revealed highly diverse and dynamic growth patterns among regions (Fig. 2), underscoring the need for a method that can robustly register developing brains at any time point to support a wide range of neurodevelopmental studies.

However, achieving whole-brain registration at every developmental stage requires repeated atlas construction, involving extensive sample collection and meticulous segmentation. Beyond the heavy workload, early postnatal stages introduce substantial variability—even among samples of the same age—making accurate registration especially challenging and prone to error. As a workaround, existing strategies^35^ often rely on interpolated atlases to fill unrepresented time points, but the underlying assumption of smooth, linear brain maturation introduces significant limitations and undermines the original goal. Brain development is inherently nonlinear and region-specific, with distinct circuits maturing on different timelines and undergoing rapid morphological changes during critical periods. Linearly transforming and averaging adjacent age templates—without accounting for these biological dynamics—risks producing anatomically inaccurate interpolated atlases, particularly in regions undergoing abrupt structural or positional shifts. Moreover, uniform transformations across the entire brain fail to capture the asynchronous development of different regions, potentially distorting key neuroanatomical features. These limitations underscore the need for data-driven atlas construction based on real samples, particularly during early postnatal stages when developmental trajectories are most divergent.

D-LMBmapX addresses this challenge by enabling automated construction of an inferred atlas for any given postnatal day from a single sample and the flanking anchor atlases, empowering the developmental neuroscience community with a foundational tool to study brain regions of interest on their own critical timelines rather than being constrained by atlas availability.

For mesoscale circuitry profiling, whole-brain axon and soma segmentation is typically the most time-consuming step. Although we previously developed D-LMBmap^17^, an automated pipeline for whole-brain circuitry profiling that substantially reduced the need for labor-intensive manual annotations, it was designed only for the adult stage. Directly extending D-LMBmap^17^ to 5D developmental analyses poses substantial challenges, as it would require extensive annotated datasets and retraining at each postnatal stage due to the marked variability in axonal and somatic morphologies throughout development. To overcome this, D-LMBmapX integrates CircuitFound, a powerful foundation model that enables large-scale self-supervised pretraining on massive brain image datasets. With only minimal sample-specific fine-tuning, this framework achieves robust and accurate axon and soma segmentation across developmental stages, offering a generalized solution for comprehensive brain mapping.

The rapid advancement of mesoscale connectomics has been driven by both large-scale 3D imaging technologies, such as LSFM, and a variety of high-resolution 2D methods^25–28^. Concurrently, there is a growing demand to integrate multi-modality, high-throughput spatial datasets that capture complementary aspects of brain organization. These include spatial transcriptomics, cross-synaptic tracing, activity-dependent labelling, and functional imaging, among others. However, due to the complexity of these procedures, such datasets are often limited to a small number of processed 2D brain slices. Existing 2D-to-3D registration methods typically require multiple consecutive sections and are constrained by limited resolution^52,53^. In this context, D-LMBmapX represents a significant advance—its ability to perform cross-modality and cross-dimensional registration allows even a single 2D section to be accurately aligned within a common 3D reference framework. This integrative capability not only enhances the scalability of mesoscale circuitry profiling but also lays the groundwork for a unified mesoscale connectomics ecosystem that incorporate anatomical, molecular, physiological, and functional data.

Forebrain TH⁺ circuits are primarily dopaminergic and noradrenergic, two critical neuromodulatory systems involved in a wide range of functions, including reward, motivation, attention, and mood regulation. Developmental deficits in these systems have long been implicated in the vulnerability to various mental disorders^54,55^. By using D-LMBmapX, we generated the first 5D whole-brain atlas of TH⁺ axonal innervation, spanning over 260 brain regions across postnatal development. While the striatum receives extremely dense innervation from P0 and cannot be segmented at the axonal level with current imaging resolution, its regional densities were quantified by intensity to ensure comprehensive coverage. In cortical areas, TH⁺ axonal density increased steadily over time, consistent with reports of prolonged growth and pathfinding of prefrontal cortex–projecting dopaminergic axons during adolescence^56^. In contrast, other regions exhibited diverse spatiotemporal dynamics, including sharp developmental surges, transient peaks (e.g., PVH and BST), or progressive declines not attributable to brain volume changes—patterns that align with evidence for system-level refinement of dopamine innervation^57^. Our atlas provides a brain-wide developmental reference for these systems and a foundation to explore how region-specific innervation trajectories relate to windows of vulnerability in neurodevelopmental and mental health conditions. The successful whole-brain profiling of TH⁺ axons highlights D-LMBmapX as an effective end-to-end workflow for mesoscale circuitry analysis, enabling precise anatomical mapping, developmental trajectory tracking, and cross-study comparisons.

D-LMBmapX marks a shift from static mapping to dynamic, integrative brain profiling, enabling automated, precise analysis of the developmental assembly of diverse neural circuits and neurodevelopmental disorders. Beyond the developing mouse brain, it offers a powerful and generalizable 5D strategy for studying growth, ageing, and disease through integrated spatial, temporal, and multi-omic analysis.

## ACKNOWLEDGEMENTS

We thank Drew Friedmann for advice on anti-TH whole brain staining. We thank the MRC Laboratory of Molecular Biology (LMB) Biological Service Group and the Ares staff for their support with animal husbandry. This work was supported by the Medical Research Council, as part of United Kingdom Research and Innovation (also known as UK Research and Innovation) [MC_UP_1201/22]. For the purpose of open access, the MRC Laboratory of Molecular Biology has applied a CC BY public copyright licence to any Author Accepted Manuscript version arising. A.M.A was supported by the MRC LMB PhD program with an MRC studentship.

## AUTHOR CONTRIBUTIONS

J.R. conceptualised the project. J.R. and Z.L. supervised the project. Z.L. led the development of D-LMBmapX. J.R. and Z.L. designed all the experiments. A.MJ.A. performed brain sample collections, tissue-clearing, immunostaining and light-sheet imaging. Z.L., P. L., Z.H., and S.Z. designed and implemented the neuron foundation model. Z. L., H.P. and J. L. designed and constructed the developmental brain atlas. Z. L., B.Z., C.L., and J.Z. designed and implemented the 2.5D registration methods. Z.L., H.P., and Y.S. designed the sample-inferred 4D atlas generation and registration pipeline. Z.L., H.P., H. L., P.L. and D.Z. designed and implemented the brain style transfer and the 2D to 3D localisation pipeline. Z.L., Y.S., and P.L. developed the D-LMBmapX software. Z.L., P.L., H.P., H.L., Z.H., Y.S., A.MJ.A, and J.R. co-annotated and checked the testing data. L.B. performed brain sample collections. R.N.S. assisted with the tissue-clearing. Z.L. and J.R. wrote the manuscript with input from all co-authors.

## DATA AND CODE AVAILABILITY

The source code of all D-LMBmapX modules, including developmental atlas construction, 2.5D registration, sample-inferred 4D atlas generation, CircuitFound, diffusion-based brain style transfer, cross-dimensional and cross-modality brain profiling, along with the executable files of D-LMBmapX software and sample data can be found at D-LMBmapX’s Github page (https://github.com/jtneuron/D-LMBmapX). Further information and requests for resources and reagents should be directed to and will be fulfilled by the lead contact, Jing Ren.

