## Supplementary figures for "D-LMBmapX: Generalised Deep Learning Pipeline for 5D Whole-brain Circuitry Profiling"

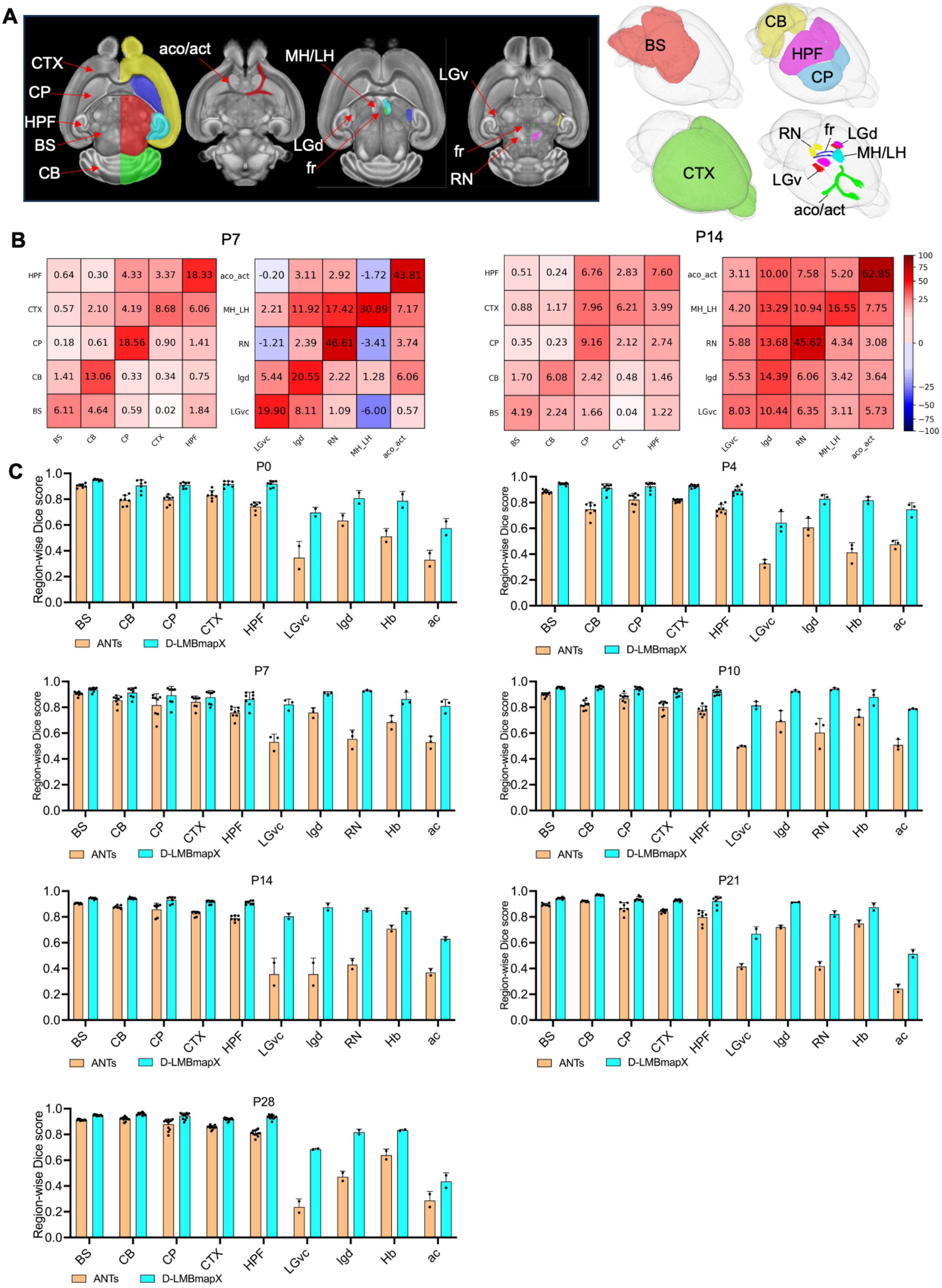


**Figure S1. Quantitative evaluation of the 2.5D registration framework with multiple constraints.**

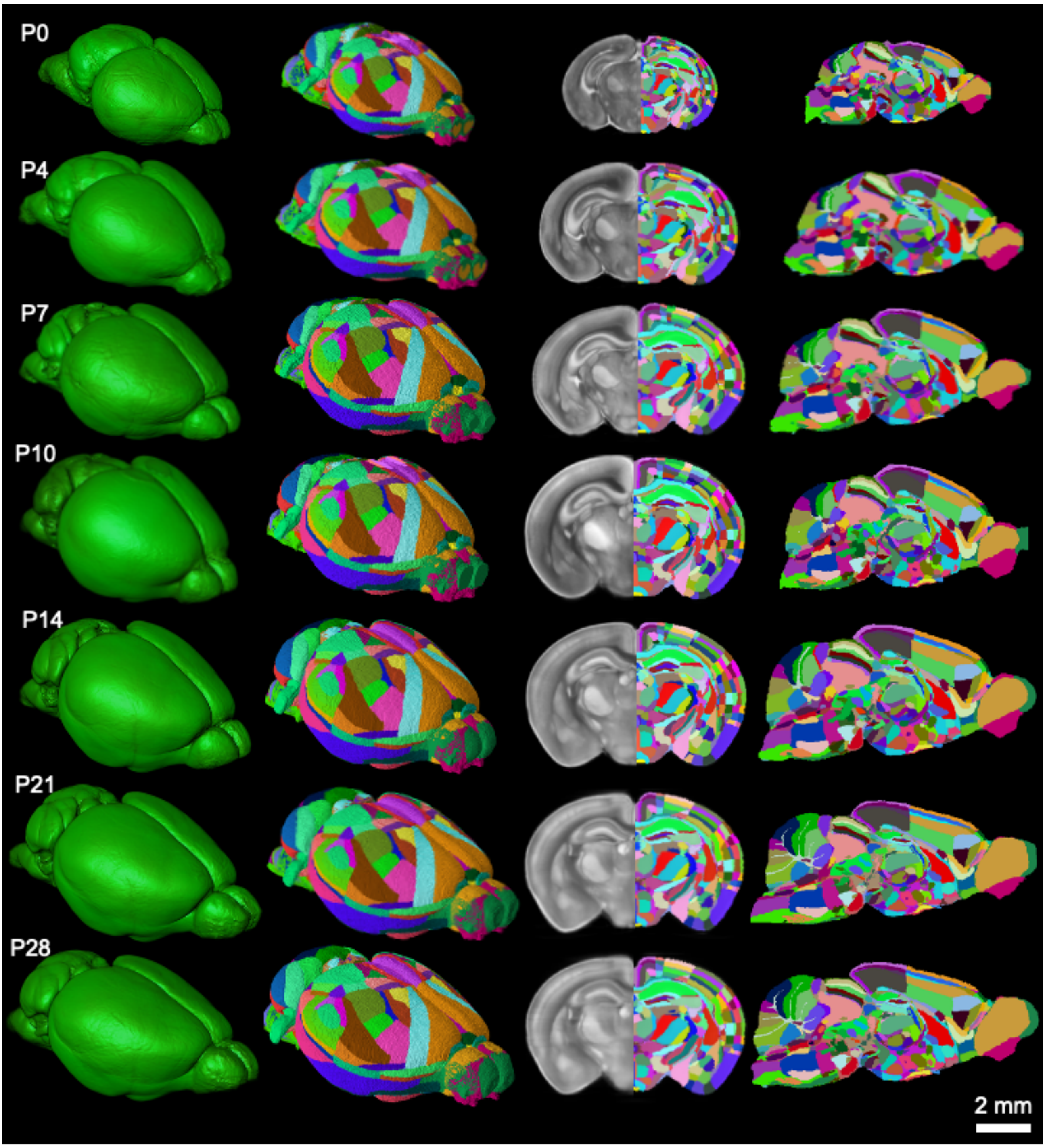


**Figure S2. Multi-view representation of the constructed 3D brain atlas across seven developmental stages.**

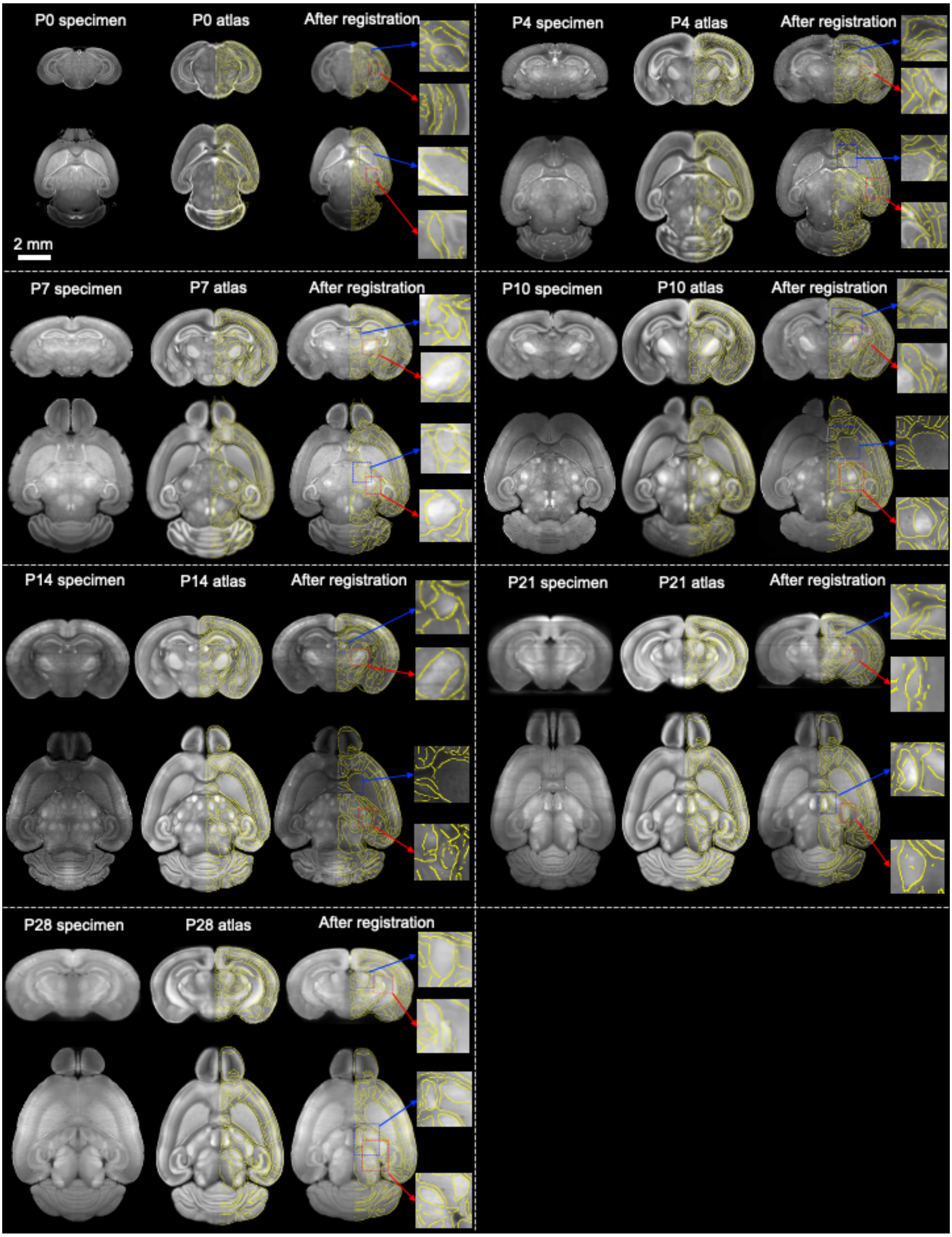


**Figure S3. Visualisation of 2.5D registration in developmental stages from P0 to P28.**

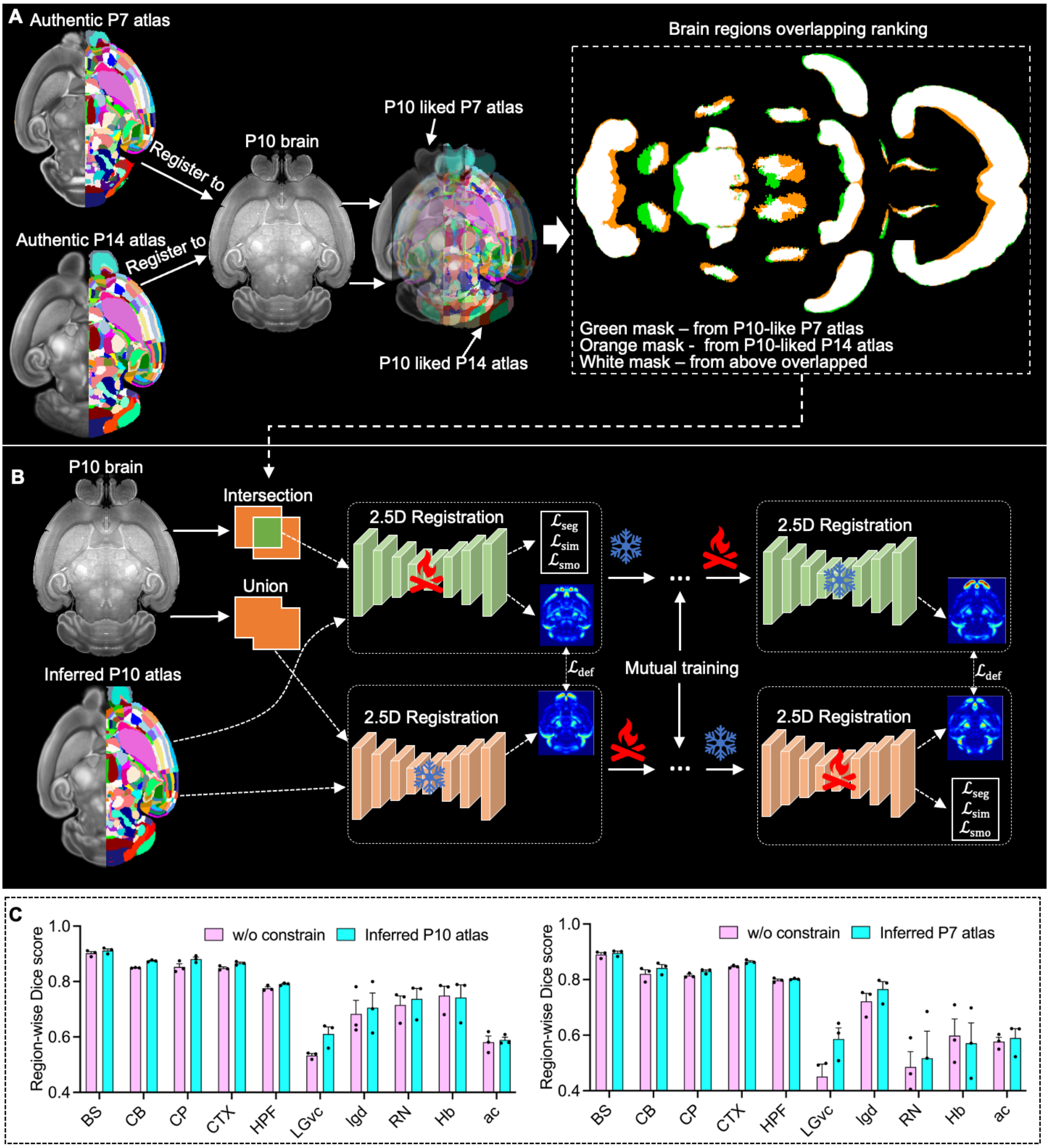


**Figure S4. Mutual learning–guided whole-brain registration using automatic flanking-atlas constraints.**

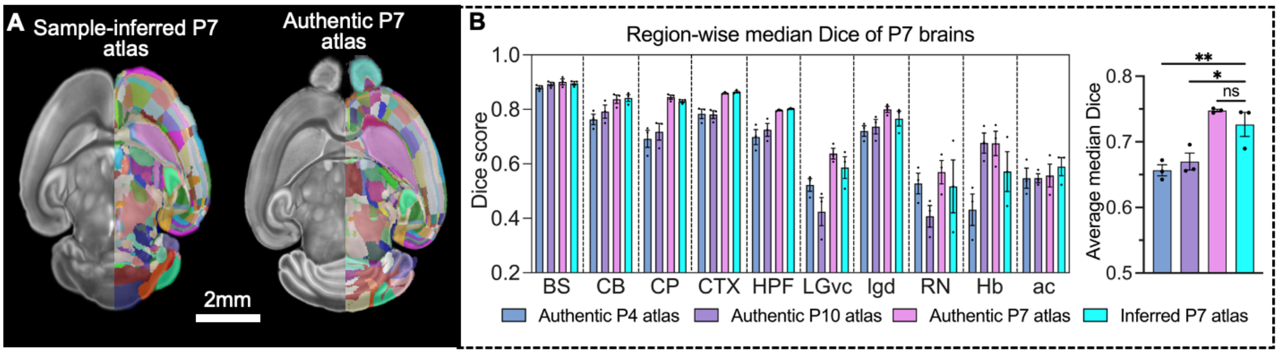


**Figure S5. Evaluation of the sample-inferred atlas at P7 in mouse brains.**

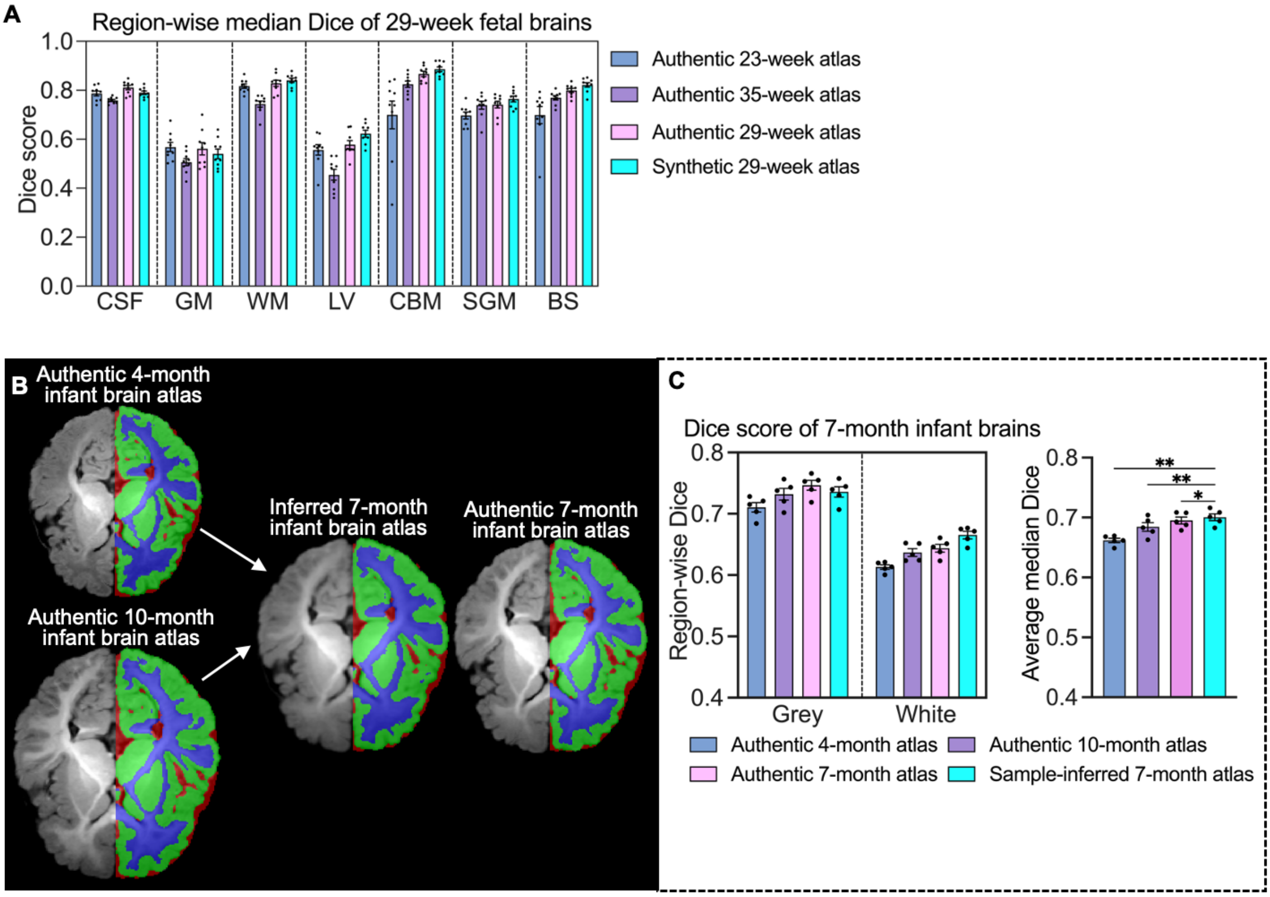


**Figure S6. Evaluation of the sample-inferred atlas construction pipeline at selected developmental stages in human fetal and infant brains.**

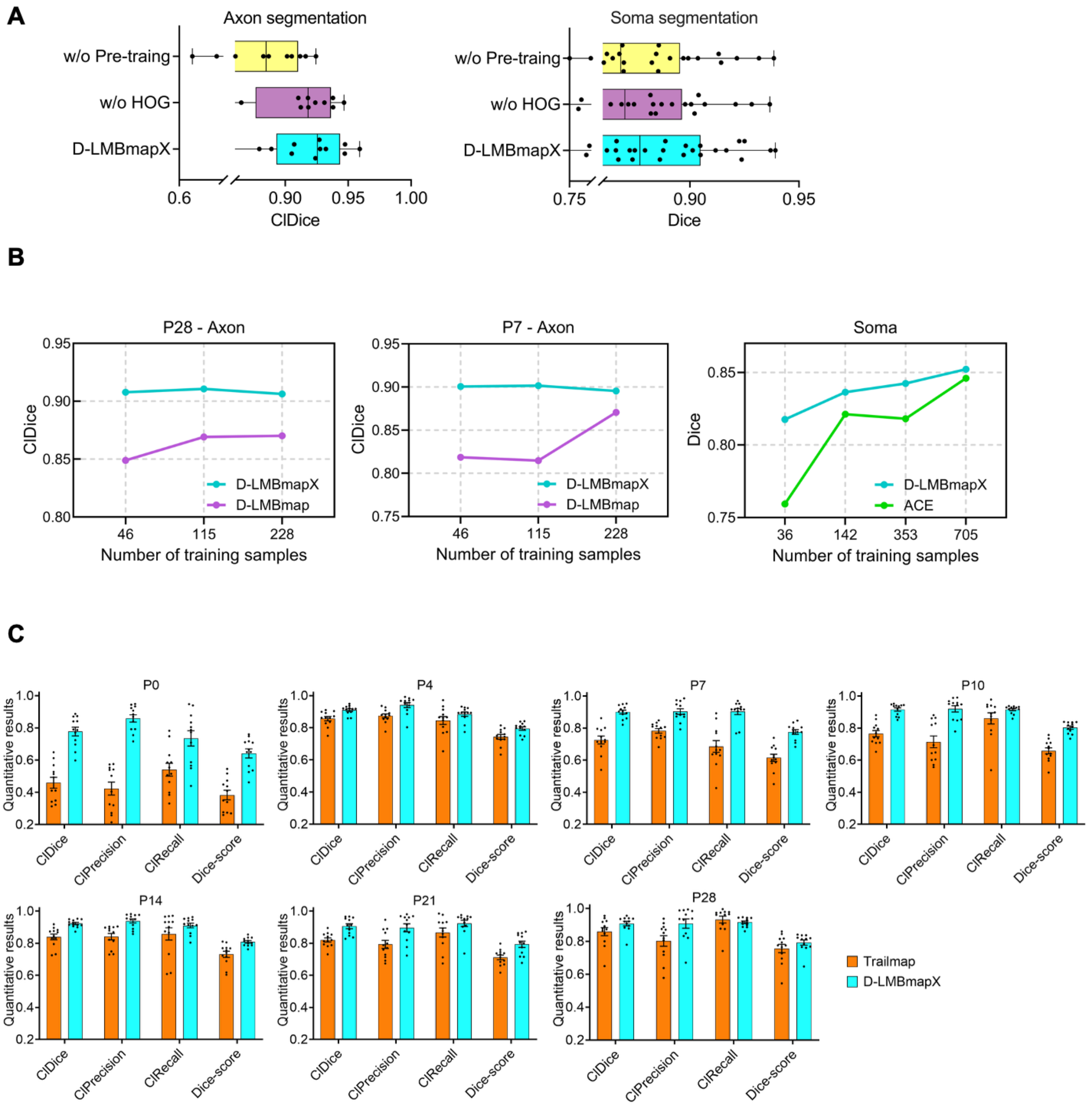


**Figure S7. Effectiveness validation of CircuitFound.**

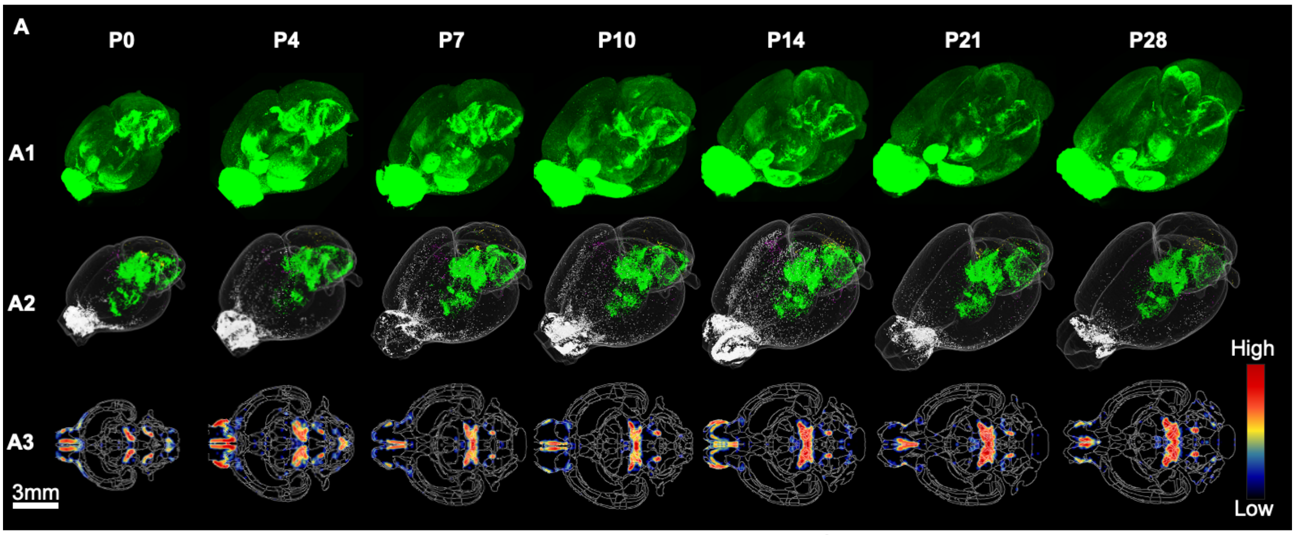

**Figure S8. Whole-brain segmentation and registration of TH⁺ somata from P0 to P28.**

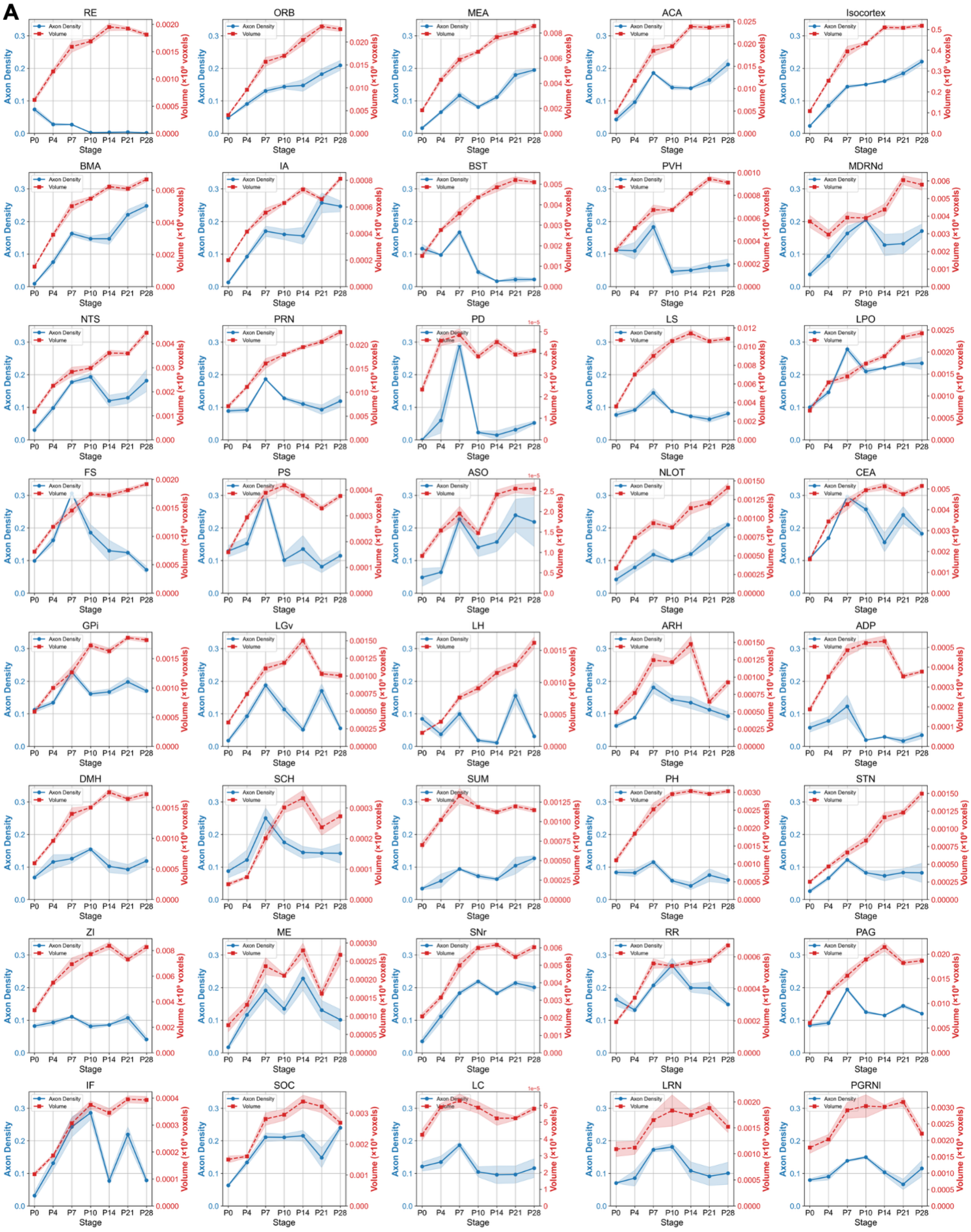


**Figure S9. Region-wise TH+ axon density changes across postnatal development.**

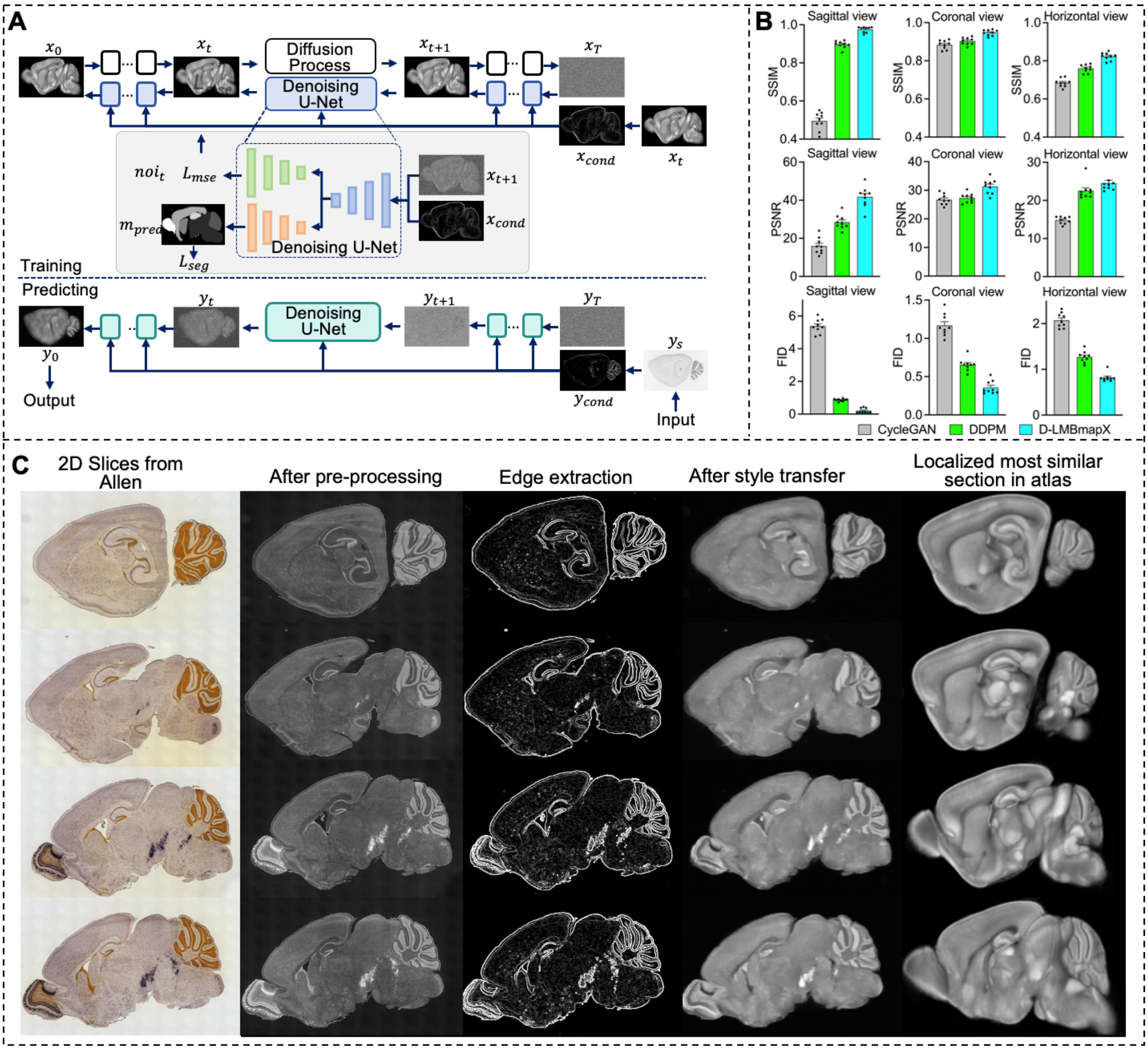


**Figure S10. Brain image style transfer based on diffusion model with brain region and edge constraints.**

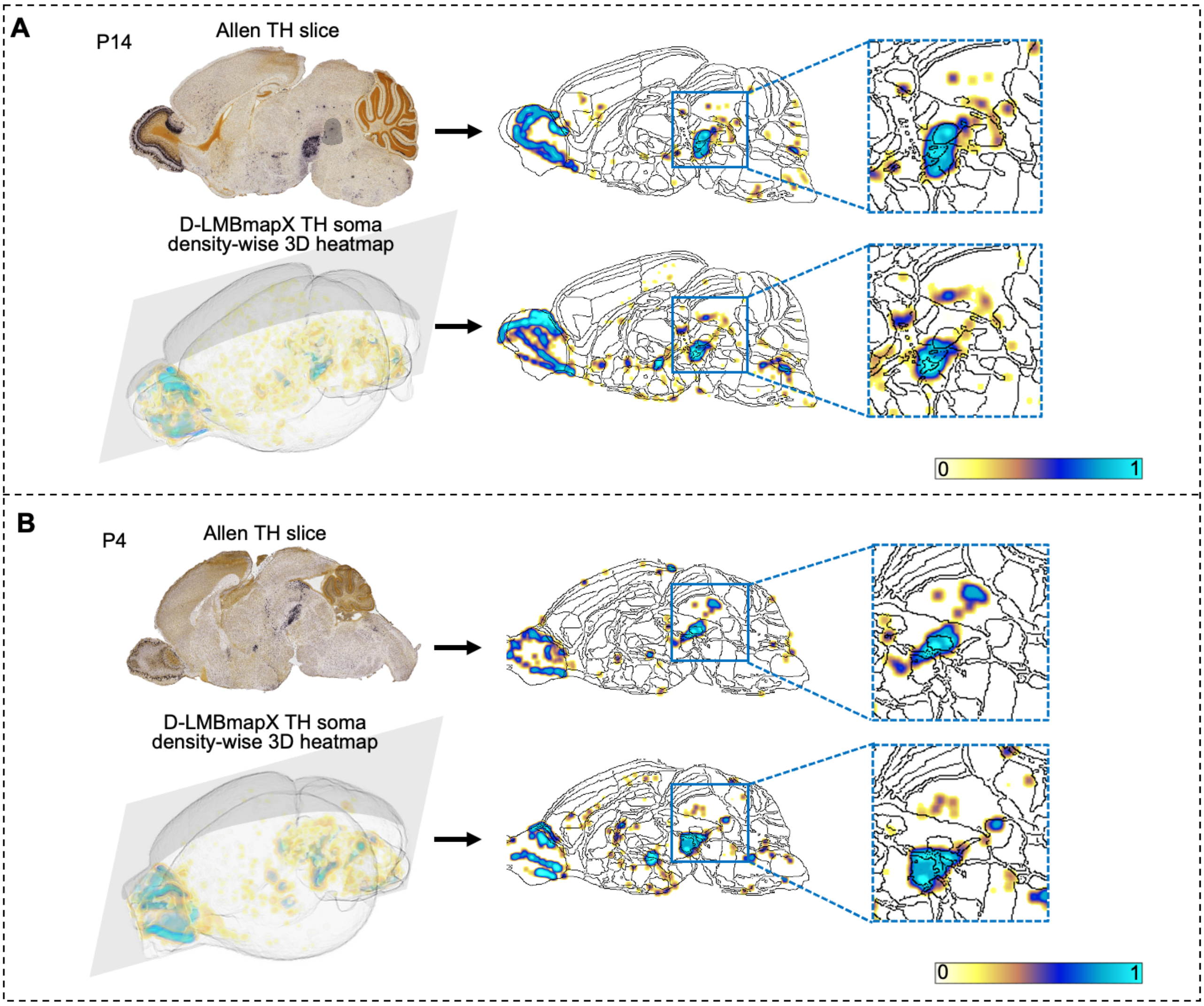


**Figure S11. Comparison of TH⁺ somata density between Allen 2D ISH slices and corresponding digital sections from 3D anti-TH–stained LSFM brains.**

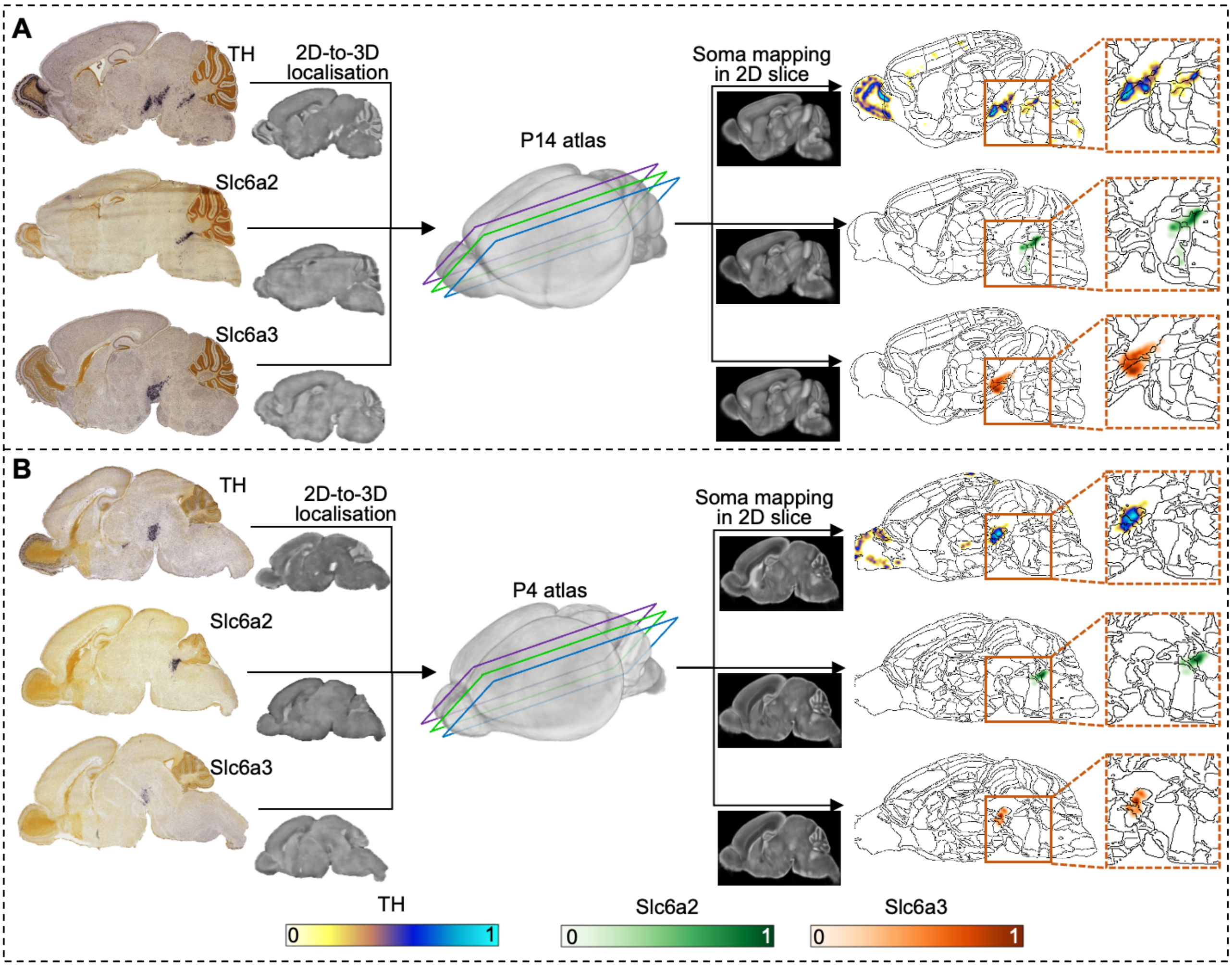


**Figure S12. Postnatal monoaminergic gene marker profiling in Allen 2D ISH slices with D-LMBmapX.**
